# Striatal and cerebellar interactions during reward-based motor performance

**DOI:** 10.1101/2025.02.06.636434

**Authors:** Joonhee Leo Lee, Agostina Casamento-Moran, Amy J Bastian, Kathleen E Cullen, Vikram S Chib

## Abstract

Goal-directed motor performance relies on the brain’s ability to distinguish between actions that lead to successful and unsuccessful outcomes. The basal ganglia (BG) and cerebellum (CBL) are integral to processing performance outcomes, yet their functional interactions remain underexplored. This study scanned participants’ brains with functional magnetic imaging (fMRI) while they performed a skilled motor task for monetary rewards, where outcomes depended on their motor performance and also probabilistic events that were not contingent on their performance. We found successful motor outcomes increased activity in the ventral striatum (VS), a functional sub-region of the BG, whereas unsuccessful motor outcomes engaged the CBL. In contrast, for probabilistic outcomes unrelated to motor performance, the BG and CBL exhibited no differences in activity between successful and unsuccessful outcomes. Dynamic causal modeling revealed that VS-to-CBL connectivity was inhibitory following successful motor outcomes, suggesting that the VS may suppress CBL error processing for correct actions. Conversely, CBL-to-VS connectivity was inhibitory after unsuccessful motor outcomes, potentially preventing reinforcement of erroneous actions. Additionally, interindividual differences in task preference, assessed by having participants choose between performing the motor task or flipping a coin for monetary rewards, were related to inhibitory VS-CBL connectivity. These findings highlight a performance-mediated functional network between the VS and CBL, modulated by motivation and subjective preferences, supporting goal-directed behavior.

## Introduction

When performing a goal-directed action, it is critical to distinguish between successful and unsuccessful outcomes that lead to reward. Previous work has implicated the basal ganglia (BG) and the cerebellum (CBL) as two subcortical systems integral for dissociating between rewarded and unrewarded motor outcomes (*1*, *2*). Despite extensive evidence showing direct anatomical projections between BG and CBL (*2–4*), there is a limited understanding of their functional connections and how they dissociate between rewarded and unrewarded motor outcomes.

The basal ganglia are a group of subcortical nuclei that process motor commands and reward information. A major functional sub-region of the basal ganglia is the ventral striatum (VS), which encompasses the nucleus accumbens and ventral parts of the putamen and is thought to be a limbic-motor interface that integrates motor and motivational signals to drive performance (*5–13*). Experimental findings point to the role of the BG in reward-based reinforcement learning (*14–16*), where actions that lead to reward are reinforced to increase the chances of future reward. In contrast, the CBL has been associated with error-based learning, adjusting motor commands following erroneous actions (*1*, *17–19*).

While it has been suggested that the VS and CBL engage independent processes for encoding rewarding and erroneous movement outcomes (*1*, *14*, *17–21*), recent experiments have shown potential communication between these regions. Anatomical tracing studies in nonhuman primates have identified direct bilateral connections between the basal ganglia and the CBL (*2*–*4*). It has also been found that the CBL encodes reward signals (*22–25*) tied to motor and non-motor outcomes and sends direct projections to the ventral tegmental area (*26*), a key input region for the VS. Human neuroimaging studies have also illustrated functional co-activation between the striatum and the CBL during motor and reinforcement learning (*27–29*). Despite converging evidence suggesting anatomical and functional links between the striatum and CBL, how they communicate when processing rewarded versus unrewarded outcomes is unclear.

Given these previous findings (*3*, *4*, *25–28*, *30*), and theoretical accounts of the links between VS and CBL (*1*, *2*, *20*), we hypothesized that these regions exhibit distinct roles in processing motor outcomes and shared roles in evaluating probabilistic, non-motor outcomes. Specifically, the VS may reinforce rewarded motor actions for motor outcomes, while the CBL processes erroneous, unrewarded motor actions, with connectivity between these regions moderated by motor performance. This interaction may facilitate the integration of reward- and error-based information to optimize motor control. For non-motor outcomes in which an individual does not have direct agency over an outcome, we hypothesized that both the VS and CBL encode reward signals in the absence of action-related errors, reflecting their broader roles in evaluating environmental outcomes that do not involve motor correction.

To test our hypothesis, we designed a skilled motor task where outcomes both depended on participants’ motor performance and events in an external probabilistic environment. Participants performed this task while their brain activity was scanned using functional magnetic resonance imaging (fMRI). In the task, participants controlled a cursor to jump through a target visualized as a variable-sized passage in a wall (**Figure 1A**). A successful motor outcome occurred when the cursor passed through the passage, while an unsuccessful motor outcome occurred if the cursor hit the wall during the jump (**Figure 1B**). An unsuccessful motor outcome resulted in immediate trial failure. Following a successful motor outcome, the task introduced an additional probabilistic event: the floor beneath the cursor’s landing point could break with varying probabilities. Importantly, this probabilistic outcome was independent of the participant’s motor performance (i.e., participants had no control over whether the floor broke). A successful probabilistic outcome occurred if the floor remained stable, whereas an unsuccessful one occurred if the floor broke. Thus, a successful trial required that two conditions were met: (1) the participant successfully controlled the cursor through the target (i.e., successful motor outcome), and (2) the floor remained stable after landing (i.e., successful probabilistic outcome) (**Figure 1B**). This task design allowed us to investigate how the ventral striatum VS and CBL respond to motor outcomes (Motor Outcome, **Figure 1A**) and how they encode probabilistic, non-motor outcomes (Probabilistic Outcome, **Figure 1A**).

**Figure 1:**
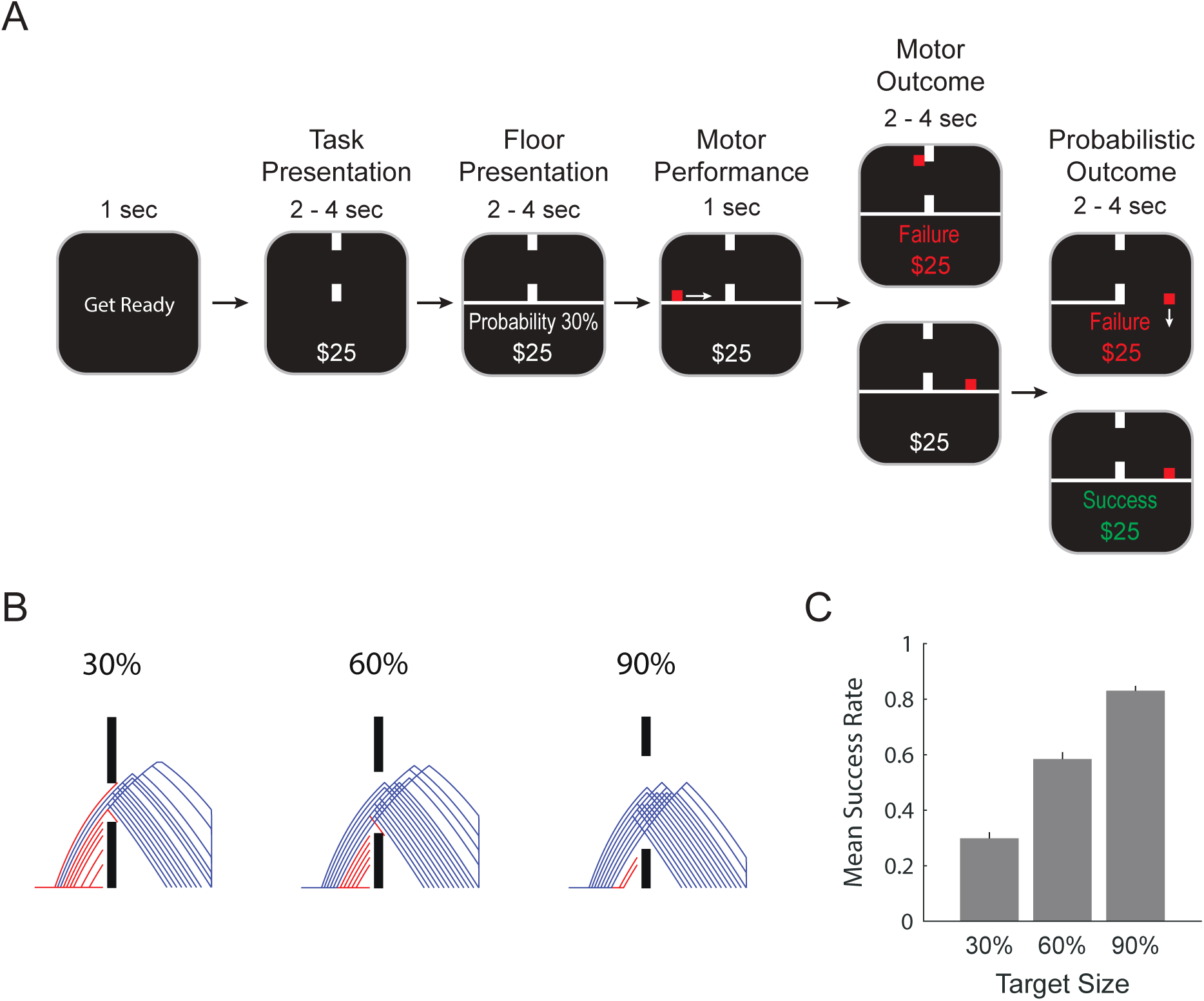
Task Design **(A)** Reward-based motor task. At the beginning of a trial, participants were presented with the target size (Task Presentation) and the probability of the floor remaining unbroken (Floor Presentation). Participants then performed the motor task by guiding the red cursor through the target (passage in the wall) and landing on the other side (Motor Performance). If the cursor hit any part of the wall, the trial failed (Unsucessful Motor Outcome, upper panel). The probabilistic outcome was revealed after the successful completion of the motor task (Successful Motor Outcome, lower panel). If the floor broke, the trial resulted in a failure, and the cursor fell off the environment (Unsuccessful Probabilistic Outcome, upper panel). If the floor remained unbroken, the trial resulted in a success (Successful Probabilistic Outcome, lower panel). **(B)** Trajectory data for an exemplary participant across the three target sizes. Blue lines indicate successful trials, and red lines indicate failed trials. Group performance (n = 28) for the three difficulty levels in the motor task. Error bars represent SEM.

## Results

We first examined differences in ventral striatum (VS) and cerebellum (CBL) activity during motor and probabilistic outcomes. Significant effects were observed in VS and CBL activity following successful and unsuccessful motor outcomes (**Figure 2**), as detailed in the following paragraphs. However, no significant differences were detected in VS or CBL activity between successful and unsuccessful probabilistic outcomes. Given this null result for probabilistic outcomes, we focused subsequent analyses on the effects observed in VS and CBL activity during motor outcomes. We present the findings from our motor outcome analyses, followed by a brief description of the probabilistic outcome results for completeness.

**Figure 2:**
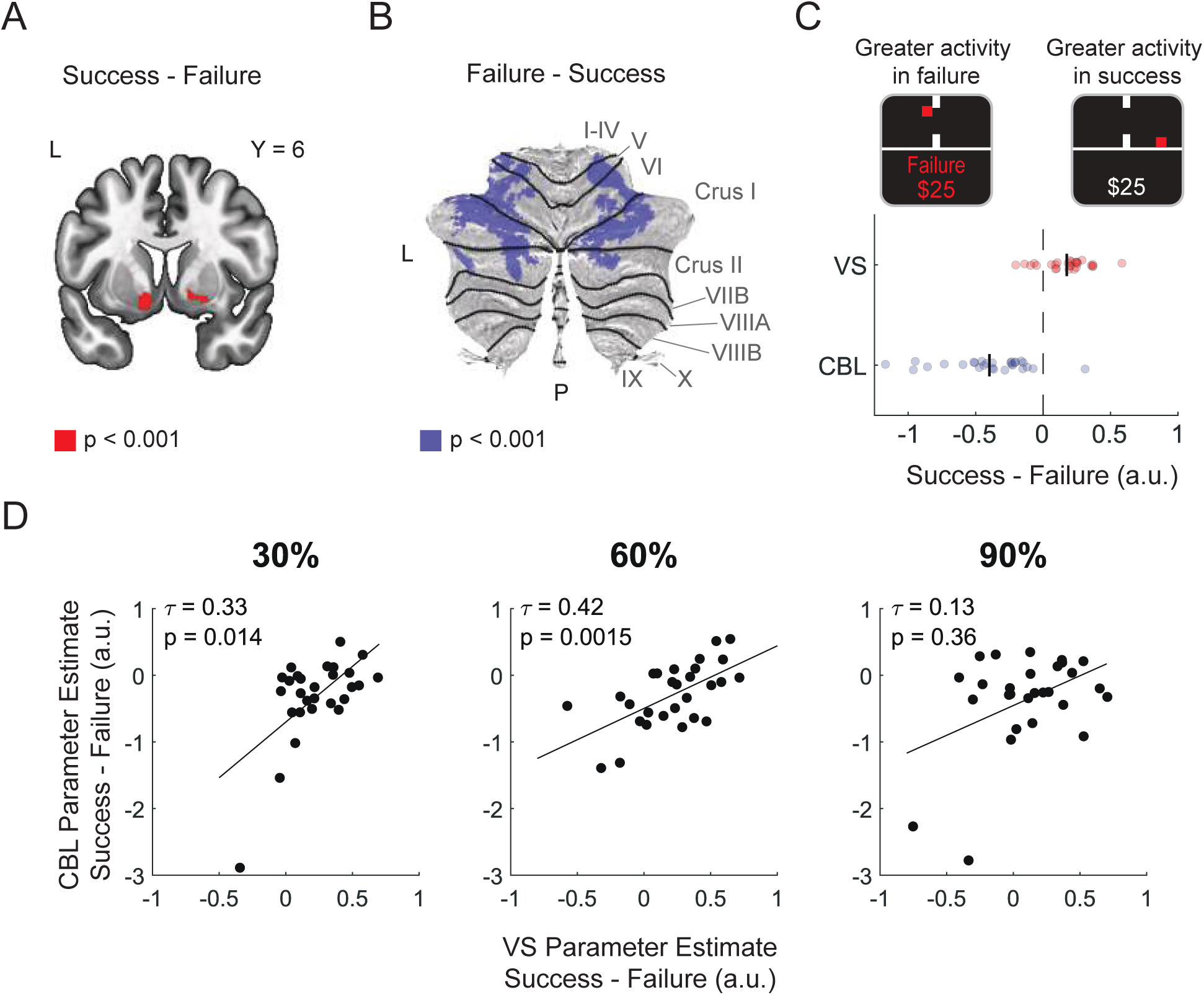
The VS and Cerebellum encode motor performance **(A)** Activity in the ventral striatum was significantly increased in successful compared to unsuccessful trials at the time of motor outcome (MNI Coordinate Peak = [-10, 6, -12], p < 0.05 FWE corrected). Contrast is displayed at p < 0.001 uncorrected. **(B)** Activity in the cerebellum (CBL) was significantly increased in unsuccessful compared to successful trials at the time of motor outcome in both anterior (MNI Coordinate Peak = [24, -36, -22], p < 0.05 FWE corrected) and posterior (MNI Coordinate Peak = [-40, -70, -22], p < 0.05 FWE corrected) regions. Contrast is displayed on a cerebellar flat map (*74*) at p < 0.001 uncorrected. **(C)** Parameter estimates for contrasts of successful versus unsuccessful motor outcomes for the VS and CBL. Each point represents a single participant. The solid vertical lines represent the mean parameter estimate for the group. VS and CBL activity at the time of motor outcome, for the three difficulty levels, across participants. We correlated the VS, and CBL parameter estimates for the Success – Failure contrast. There was a significant positive correlation between the VS and CBL activity for the hard (30%, Kendall’s τ = 0.33, p = 0.014) and medium (60%, Kendall’s τ = 0.42, p = 0.0015) target sizes. There was a non-significant positive trend observed for the easy target size (90%, Kendall’s τ = 0.13, p < 0.36). These results indicate that for individu- als with greater VS activity following successful motor performance, the CBL exhibited reduced activity following failed motor performance.

### The VS and Cerebellum encode successful and unsuccessful motor outcomes

We examined individuals’ performance at different difficulty levels during the incentivized skilled motor task. Participants’ mean success rate for the easy, medium, and hard difficulties was approximately 30%, 60%, and 80%, respectively (**Figure 1B, C**), which matched the thresholded target probabilities (i.e., 30%, 60%, and 90%). Participants’ motor performance was significantly modulated by target size (mixed-effects linear model, t_(1202)_ = 31.2, p < 0.001) (**Supplementary** Figure 2). While session number was a significant predictor of motor performance (mixed-effects linear model, t_(1202)_ = 4.632, p < 0.001), these effects were small (∼2% increase per session) (**Supplementary** Figure 2). The probabilistic floor outcome was not a significant predictor of performance (t_(1202)_ = 0.6273, p = 0.53), indicating that participants’ performance in the motor task was not influenced by the probability of the floor remaining unbroken following successful motor execution (**Supplementary** Figure 2B). These results illustrate that participants’ performance remained relatively stable throughout the incentivized phase of the experiment and that the thresholded target size matched the intended performance probabilities. This allowed us to independently investigate activity in the VS and CBL relative to motor and probabilistic outcomes, controlling for potential learning effects and cross-talk between participants’ motor performance and probabilistic, non-motor outcomes.

We compared conditions following successful and unsuccessful motor outcomes (Motor Outcome, **Figure 1A**) to determine how the ventral striatum (VS) and cerebellum (CBL) respond. Activity in the bilateral VS was more active following successful compared to unsuccessful motor outcomes (MNI Coordinate Peak = [-10, 6, -12], p < 0.05 _FWE corrected_; **Figure 2A, C**), and activity in both the anterior (MNI Coordinate Peak = [24, -36, -22], p < 0.05 _FWE corrected_) and posterior (MNI Coordinate Peak = [-40, -70, -22], p < 0.05 _FWE corrected_) regions of the CBL was more active following unsuccessful compared to successful motor outcomes (**Figure 2B, C**). At an uncorrected threshold of p < 0.001, activity in the anterior CBL spanned lobules V-VI, and activity in the posterior CBL spanned Crus I and II (**Supplementary Table 5**). These results align with our initial hypothesis and previous studies suggesting that the VS encodes successful motor outcomes (*1*, *14*, *27*), whereas the cerebellum is sensitive to motor errors in unsuccessful motor outcomes (*1*, *17–19*).

Next, we examined differences in activity in VS and CBL during successful and unsuccessful probabilistic outcomes (Probabilistic Outcome, **Figure 1A**). Based on our initial hypothesis, we expected both the VS and CBL to encode reward signals, exhibiting greater activity during successful probabilistic outcomes than unsuccessful ones. However, at the specified statistical threshold of p < 0.05 _FWE corrected_, we did not observe any significant differences in activity between successful and unsuccessful probabilistic outcomes in either the VS or the CBL. With this in mind, we focused our subsequent analyses on the time of motor outcome.

While the VS and CBL have been considered distinct subcortical systems with separate roles in encoding reward and error-based outcomes, recent studies have revealed direct anatomical connections between these regions (*2–4*, *30*). To investigate how signals in VS and CBL at the time of motor outcome are functionally related, we examined how the relationship between activity in these regions was influenced by expectations of motor performance (i.e., difficulty level) (**Figure 2D**). We observed significant positive correlations between the VS and CBL parameter estimates, comparing successful and unsuccessful motor outcomes for the hard (30%, Kendall’s τ = 0.33, p = 0.014) and medium (60%, Kendall’s τ = 0.42, p = 0.0015) target sizes. A non-significant but positive trend was observed for the easy target size (90%, Kendall’s τ = 0.13, p = 0.36), which could be attributed to the high success rate and reduced sampling of unsuccessful motor outcomes. Together, these results indicate that individuals with greater VS activity following successful motor outcomes had reduced CBL activity following unsuccessful motor outcomes. This supports the idea that activity in the VS and cerebellum have complementary roles in representing motor outcomes.

### Successful and unsuccessful motor outcomes modulate VS-CBL connectivity

To further test our hypothesis that the VS and CBL communicate to encode motor outcomes, we constructed a dynamic causal model (DCM) to assess specific connectivity profiles between these two regions. We specified a deterministic, bilinear DCM to infer directed influences between the VS and CBL (i.e., effective connectivity) and the modulation of this connectivity by successful or unsuccessful motor outcomes (**Figure 3A**). Importantly, the sign of the connections indicates whether a region’s activity is enhanced (positive/excitatory) or suppressed (negative/inhibitory) due to hypothesized direct experimental inputs or inputs from other brain regions (*31–33*). We constructed a fully connected DCM for each participant that enabled all motor outcomes (Motor Success & Motor Failure) to drive activity in VS and CBL and modulate connections between the VS and CBL. We then used parametric empirical Bayes (PEB) analyses to formally test how successful or unsuccessful motor outcomes could influence the connectivity between and within the VS and CBL. This process involves comparing the full model to reduced candidate models with certain combinations of connections (**see methods**).

**Figure 3:**
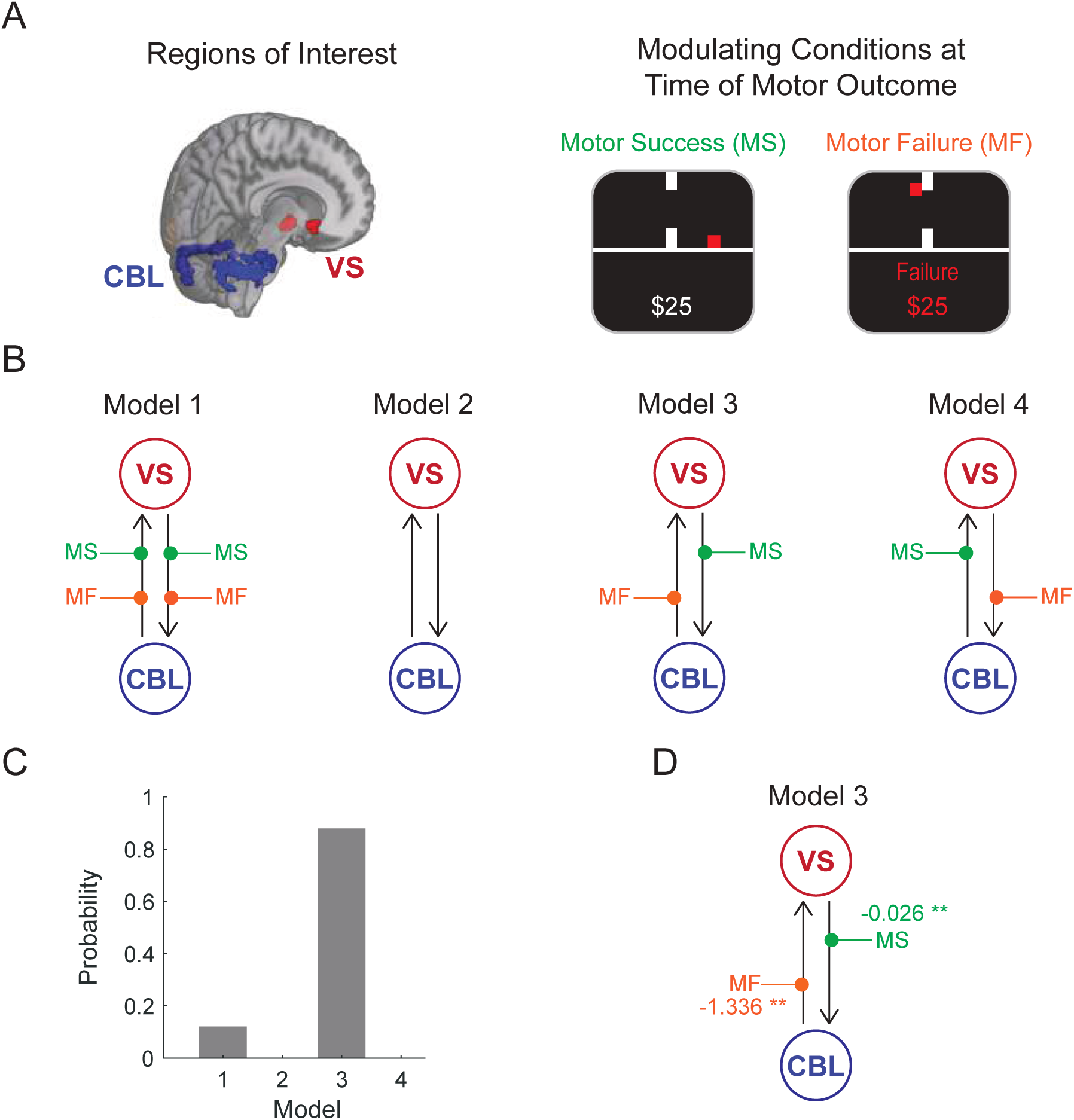
Performance outcomes modulate VS-CBL connectivity **(A)** Regions of interest and modulating conditions for the DCM analysis. Regions were the VS (red) and CBL (blue), and the modulating conditions were successful (MS, green) and unsuccessful (MF, orange) motor outcomes. VS activity was extracted using an a priori mask and the CBL activity was extracted using a leave-one-subject-out (LOSO) ROI based on the contrast in Figure 2C. **(B)** DCM model specification. We specified four potential models that varied according to how successful (MS) and unsuccessful (MF) motor outcomes modulated VS-CBL effective connectivity (black arrows). **(C)** A Bayesian model comparison showed that Model 3 was the winning model (88%) for describing the commonalities (average connectivity) across participants’ VS-CBL connectivity. **(D)** Bayesian Model Average (BMA) of Parametric Empirical Bayes (PEB) parameters highlighting the impact of successful and failed motor outcomes on VS-CBL connectivity. The VS inhibits CBL following successful motor outcomes, whereas the CBL inhibits VS following unsuccessful motor outcomes. **indicates that PEB parameters have very strong positive Bayesian evidence (posterior probability > 0.99).

We first investigated whether successful and unsuccessful motor outcomes directly influenced activity in the VS and CBL by defining four candidate models (**Supplementary** Figure 3A). Model 1 (full model) enabled all motor outcomes to drive VS and CBL activity, whereas Model 2 (null model) did not allow any driving inputs. Model 3 was designed based on previous studies showing structural and functional connections between VS and CBL and our GLM results, where the VS would receive direct inputs following successful motor outcomes (representative of reward receipt), and the CBL would receive direct inputs following unsuccessful motor outcomes (representative of motor error). Finally, Model 4 was specified as the inverse of Model 3. A Bayesian model comparison revealed that Model 1 best explained the influence of motor outcomes on activity in VS and CBL (90%, **Supplementary** Figure 3B). To further examine the connections of each parameter, we computed the Bayesian Model Average (BMA), which averaged parameters from different models weighted by each model’s posterior probabilities (*34*, *35*). We observed positive evidence for each direct input (**Supplementary** Figure 3C), suggesting that successful and unsuccessful motor outcomes directly modulate the VS and CBL. Based on the model comparison and BMA results, we decided to enable motor outcomes to modulate both the VS and CBL when investigating the modulatory connections between the regions.

To test how successful and unsuccessful motor outcomes modulate VS-CBL connectivity, we constructed four candidate models (**Figure 3B**) that varied in their modulatory connections. Model 1 (full model) enabled all modulatory connections, whereas Model 2 (null model) disabled all modulatory connections. Model 3 specified that successful motor outcomes modulate the connectivity from the VS to the CBL, and unsuccessful motor outcomes modulate the connectivity from the CBL to the VS. Finally, Model 4 was created as the inverse of Model 3. A Bayesian model comparison revealed that Model 3 (88%) best explained how motor outcomes modulated the VS- CBL connectivity (**Figure 3C**). By performing BMA, we observed very strong evidence of inhibition from the VS to the CBL (-0.026, p > 0.99) following successful motor outcomes and from the CBL to the VS (-1.336, p > 0.99) following unsuccessful motor outcomes (**Figure 3D**). These results are consistent with the idea that CBL encodes motor errors, which may inhibit reinforcing VS activity during unsuccessful motor outcomes. In the case of successful performance, VS may inhibit CB activity since motor error signals are not necessary to update actions that lead to a successful motor outcome.

### VS-CBL connectivity predicts participants’ subjective preferences for motor performance

Recent studies have indicated that prospective incentives influence motor performance, and differences in connectivity within the motor system capture the degree to which individuals respond to such incentives (*9*, *12*, *36*, *37*). In particular, the VS is considered a limbic-motor interface that integrates motoric signals with various factors, such as motivation or subjective preferences for potential rewards (*5–13*). As the CBL is another key region for updating and processing motor signals, we investigated whether differences in motor performance and subjective preferences for the motor task influenced VS-CBL connectivity. We measured participants’ motor performance by calculating their mean success rate in the motor task, and inferred participants’ subjective preferences for performing the motor task by asking them to choose between performing the motor task or accepting a probabilistic outcome (i.e., coin flip) for potential rewards (**Figure 4A**, see methods). We defined participants’ subjective preference for the motor task as the mean acceptance rate for choosing the motor task over the probabilistic outcome.

**Figure 4:**
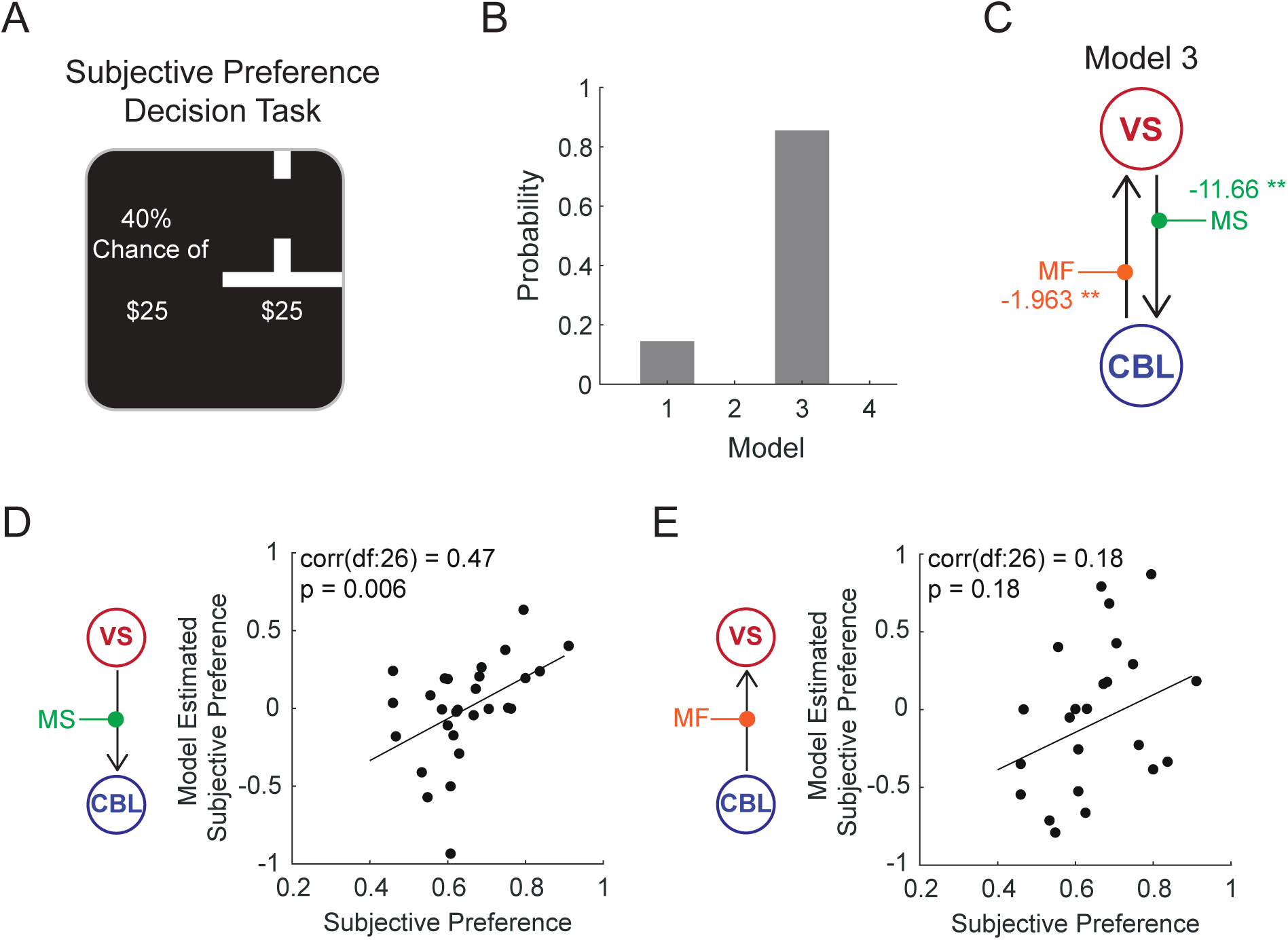
VS-CBL connectivity predicts participants’ preference for performing the motor task **(A)** Prior to performing the reward-based motor task in the scanner, participants made a series of choices to either perform motor tasks of varying difficulty or accept potential probabilistic outcomes (weighted coin flip) for a potential monetary reward of $25. The target size associated with the motor task was sampled from the three target sizes used in the reward-based motor task (Figure 1 B, C). The coin flip had a proba- bility of success varying from 0.10 to 0.90. Participants’ subjective preference was computed as the mean acceptance rate of the motor task over the coin flip. **(B)** A Bayesian model comparison showed that Model 3 was the winning model (85% for describing the effect of participants’ subjective preference for performing the motor task on their VS-CBL connectivity. **(C)** Bayesian Model Average of Parametric Empirical Bayes (PEB) parameters illustrating the impact of participants’ subjective preference for performing the motor task on VS-CBL connectivity. Individuals with a greater preference to perform the task exhibited increased inhibitory connections from the VS to the CBL following successful motor outcomes and from the CBL to the VS following failed motor outcomes. ** indicates that PEB parameters have very strong positive Bayesian evidence (posterior probability > 0.99). **(D)** Out-of-sample estimation of participants’ subjective preferences for performing the task using the degree of inhibition of the CBL by the VS following successful motor outcomes (PEB parameter, Left). Out-of-sample estimation of participants’ subjective preference using the degree of inhibition of the VS by the CBL following failed motor outcomes (PEB parameter, Left).

We first investigated which of the four candidate models (**Figure 3B**) captured the modulation of VS-CBL by motor outcomes best explained interindividual differences in mean motor performance. We found that no model explained interindividual differences in mean motor performance better than chance, with model 2 (null model) showing the highest probability (44%) (**Supplementary** Figure 4A). In addition, there was no positive evidence of mean motor performance modulating the effect of motor outcomes on VS-CBL connectivity (**Supplementary** Figure 4B). Thus, there was insufficient evidence to suggest that participants’ performance in the motor task was influenced by interindividual differences in VS-CBL connectivity.

Next, we investigated which of the four candidate models best explained interindividual differences in subjective preference for the motor task. Model 3 (85%) best explained interindividual differences in subjective preferences. By performing BMA, we observed very strong evidence that individuals with greater subjective preferences for performing the motor task showed greater inhibitory connections from the VS to CBL (-11.66, p > 0.99) following successful motor outcomes and greater inhibitory connections from the CBL to VS (-1.963, p > 0.99) following unsuccessful motor outcomes (**Figure 4C**). Overall, our results indicate that the VS and CBL inhibit one another following successful and unsuccessful performance. In addition, these inhibitory connections are strengthened for individuals who display a greater preference for performing the motor task for potential reward. These results suggest that the dissociable roles of VS and CBL in processing incentivized performance, and motor errors are further amplified in individuals with greater preferences for performing motor tasks.

To verify and validate the associations between the VS-CBL connectivity and subjective preferences, we performed a series of leave-one-out (LOO) cross-validation analyses. In these analyses, a PEB model was fitted to all but one participant, and the covariate (i.e., subjective preference for performing the motor task) for that left-out participant was predicted. Importantly, we repeated this procedure twice where the PEB parameter used to predict the covariate was 1) the modulation from VS to CBL following successful motor outcomes (**Figure 4D**, left) and 2), the modulation from CBL to VS following unsuccessful motor outcomes (**Figure 4E**, left). We observed significant correlations between participants’ predicted and actual subjective preferences when the predicted values were estimated using the modulation from VS to CBL following successful motor outcomes (r(_26_) = 0.47, p = 0.006, **Figure 4D**, right), but not when using the modulation from CBL to VS following unsuccessful motor outcomes (r(_26_) = 0.18, p = 0.18, **Figure 4E**, right). From these results, the effect size associated with the inhibitory modulation from VS to CBL following successful motor outcomes predicted individuals’ subjective preferences for the motor task.

### Motor outcomes modulate VS-CBL connectivity for both the sensorimotor and cognitive CBL

The CBL activations we see in our GLM analysis (**Figure 2C**), and the CBL ROI we specified for the first DCM analysis spanned both the anterior and posterior regions of the CBL. The anterior CBL (lobules I-V) and lobule VI are thought to process sensorimotor information (*38–40*), while the posterior CBL (predominantly Crus I & II) is involved in more cognitive processes (*39–41*) such as working memory (*42*), social cognition (*43*), and reward encoding (*22–25*). Thus, we investigated whether the VS-CBL connectivity during motor outcomes differed between the sensorimotor (Mtr. CBL) and cognitive (Cog. CBL) cerebellar regions. We first specified ROIs corresponding to the Mtr. and Cog. CBL using the leave-one-subject-out (LOSO) approach (**see methods**). Based on our GLM results (**Figure 2B**), the Mtr. CBL ROI included activations from lobules IV-VI while the Cog. CBL ROI included activations from Crus I and II (**Figure 5A**). We then specified a deterministic, bilinear DCM that resembled the initial DCM analysis (**Figure 3**), with the key difference that the model allowed for directed influences between the VS-Mtr. CBL and VS-Cog. CBL connections (**Figure 5**). In addition, we switched off the connections between the Mtr. and Cog. CBL as we were primarily interested in connections between the VS and CBL. Next, we defined a set of candidate models to identify the best explanation for connectivity between the VS and the Mtr./Cog. CBL following motor outcomes. The first factor varied how successful or unsuccessful motor outcomes modulated VS-CBL connectivity (**Figure 5B**, up). The second factor varied in terms of whether motor outcomes modulated the VS-Mtr. CBL connectivity or VS-Cog. CBL connectivity (**Figure 5B**, down). Each factor consisted of three levels, and with the addition of a null model (no modulations), we specified a total of 3 x 3 + 1 = 10 candidate models (**Supplementary** Figure 5).

**Figure 5:**
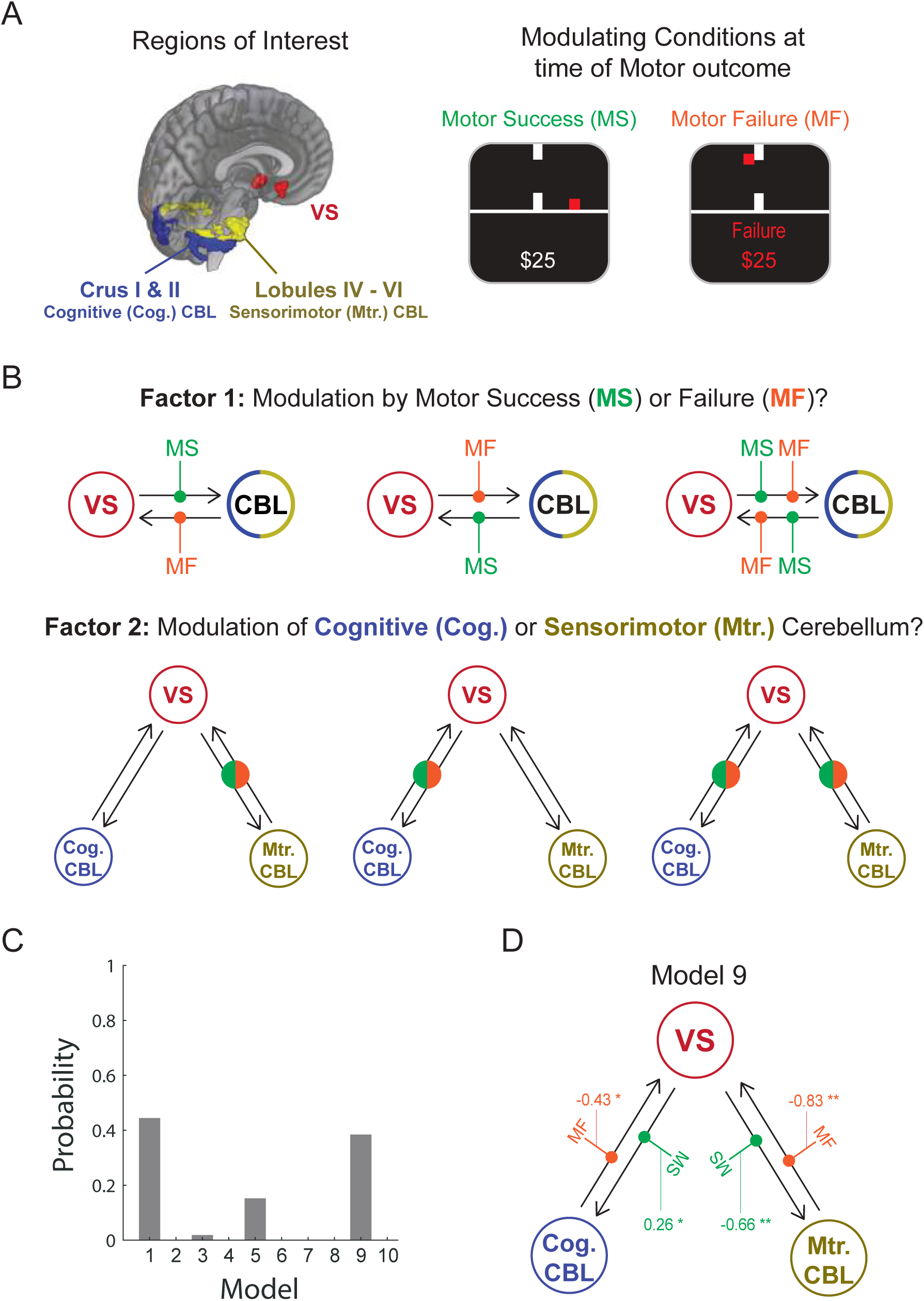
Motor outcomes modulates VS-CBL connectivity for both the sensorimotor and cognitive CBL. **(A)** Regions of interest and modulating conditions for the DCM analysis were specified in a similar fashion as the original DCM analysis (Figure 3A), with the exception that the CBL was divided into the sensorimotor (Mtr. CBL) and cognitive (Cog. CBL) subregions. The Mtr. CBL consisted of activations in lobules IV, V and VI, whereas the Cog. CBL consisted of Crus I and II. **(B)** Candidate PEB models were specified according to the two factors. The open blue-yellow circles indicate that factor 1 (modulation by motor success/failure) did not consider the specificity of the Mtr. (yellow) or Cog. CBL (blue). The filled green-orange circle indicates Factor 2 (modulation of Mtr. or Cog. CBL) did not consider the modulation of regions due to task success (green) or failure (orange). **(C)** A Bayesian model comparison showed that models 1 (44%) and 9 (38%) best described the commonalities (average connectivity) across participants’ VS-Mtr. CBL and VS-Cog. CBL connectivity. The VS inhibits Mtr. CBL and excites Cog. CBL following successful motor outcomes. The Mtr. and Cog. CBL inhibits VS following failed motor outcomes. ** PEB parameters showing very strong positive Bayesian evidence (posterior probability > 0.99). * indicates that PEB parameters show positive Bayesian evidence (posterior probability > 0.73).

We found that no model explained the VS-Mtr./Cog. CBL connectivity with greater than 50% of the model evidence, with Model 1 (full model with all connections, 44%) and Model 9 (38%) having the two greatest probabilities (**Figure 5C**). Model 9 allowed VS to modulate both the Mtr. and Cog. CBL following successful motor outcomes and for the Mtr./Cog. CBL to modulate the VS following unsuccessful motor outcomes (**Figure 5D**). As there was no definitive winning model, we performed BMA to obtain averaged estimates of each connection. We found very strong evidence of the VS inhibiting the Mtr. CBL (-0.66, p > 0.99) following successful motor outcomes and the Mtr. CBL inhibiting the VS (-0.83, p > 0.99) following unsuccessful motor outcomes (**Figure 5D**, **Supplementary Table 2**). Interestingly, we found positive evidence of the VS exciting the Cog. CBL (0.26, p > 0.73) following successful motor outcomes and the Cog. CBL inhibiting the VS (-0.43, p > 0.73) following unsuccessful motor outcomes (**Figure 5D**, **Supplementary Table 2**). These results suggest that the VS-Mtr. CBL primarily drives the inhibitory connections between the VS and CBL following motor outcomes. In addition, our results indicate that the VS could differentially communicate with the Mtr. and Cog. CBL.

Finally, we investigated whether interindividual differences in subjective preferences for the motor task can be explained by VS-Mtr. CBL or VS-Cog. CBL connectivity. By repeating the analysis in Figure 4 with this second DCM (see methods), we found evidence suggesting that individuals with greater subjective preference for the motor task showed greater inhibitory connections from the VS to Cog. CBL (-4.86, p > 0.73) following successful motor outcomes and greater inhibitory connections from the Cog. CBL to VS (-5.091, p > 0.73) following unsuccessful motor outcomes (**Supplementary Table 2**). Importantly, we failed to find a significant relationship between subjective preferences and modulation of VS-Mtr. CBL connectivity following motor outcomes (p < 0.73, **Supplementary Table 2**). By performing the LOO cross-validation analysis, we found that predicted subjective preferences for the motor task estimated using the modulation from VS to Cog. CBL (following successful motor outcomes) were significantly correlated with individuals’ actual subjective preferences (r(_26_) = 0.32, p = 0.047, **Supplementary** Figure 6B). Thus, modulation from VS to Cog. CBL following successful motor outcomes significantly predicted individuals’ subjective preferences for the motor task. These findings suggest that there is a transition from excitation of the Cog. CBL by the VS to inhibition following successful motor outcomes with increased subjective preferences. Thus, subjective preferences could modulate the VS-Cog. CBL connectivity, which aligns with recent studies suggesting that the Cog. CBL plays a role in reward processing (*22–26*).

## Discussion

Although many studies have shown anatomical and functional coupling between the VS and CBL (*2–4*, *28*, *29*, *44*, *45*), and both regions play integral roles in motor learning and reward processing (*1*, *5*, *9*, *14*, *24*, *25*, *46*, *47*), the functional mechanisms underlying how the VS and CBL interact when tying motor actions with rewarded and unrewarded outcomes are unclear. Here, we demonstrate that the VS and CBL dissociate between successful and unsuccessful motor outcomes and play complementary roles in reward-based error encoding. Importantly, we find that the VS inhibits the CBL following successful motor outcomes, whereas the CBL inhibits the VS following unsuccessful motor outcomes. These modulatory connections are stronger for individuals with a stronger preference for performing the motor task. Our findings go beyond previous studies that have primarily focused on anatomical connections or isolated roles of CBL or VS in reward and motor performance - we show that the VS forms different modulatory connections with the sensorimotor and cognitive CBL when integrating motor outcomes with reward information.

Our findings support the idea that the VS and CBL underly the roles of reward-based reinforcement and error-driven processes, respectively (*1, 20*). We observe stronger activity in the VS following successful versus unsuccessful motor outcomes, which aligns with several studies showing that the striatum encodes rewarding outcomes (*5*, *14*, *16*, *27*, *48*, *49*). On the other hand, the CBL was more active for unsuccessful versus successful motor outcomes, consistent with the role of CBL in error processing (*1*, *17*, *18*). Prior studies have found neural correlates of error processing in the CBL in reaching movement tasks (*50–52*). However, these studies mainly showed activations in the anterior, sensorimotor CBL regions and did not find significant activations in the posterior regions such as Crus I & II. Unlike these previous studies, our motor task was tied to prospective monetary rewards, which may explain why we observe activations in Crus I & II as recent studies have suggested the involvement of these regions in reward processing (*22–25*).

Our data provides new insight into the role of the connectivity between the VS and CBL following motor outcomes. Across individuals, we observe that those with more ventral striatal sensitivity to reward (successful motor outcomes) exhibit reduced error-related (unsuccessful motor outcomes) signals in the CBL and vice versa, suggesting inhibitory VS-CBL connections. This hypothesis was supported in our DCM analysis, where the VS inhibited the CBL following successful motor outcomes, and the CBL inhibited the VS following unsuccessful motor outcomes. These findings suggest that following successful motor outcomes, the VS could inform the CBL not to process error signals; while following unsuccessful motor outcomes, the CBL could inform the VS not to reinforce erroneous motor actions. Our model comparison focusing on direct inputs revealed that the best model enabled successful and unsuccessful motor outcomes to modulate both VS and CBL. Thus, while the VS and CBL could potentially process all motor outcomes, communication between both regions may enable them to distinguish between rewarding and erroneous performance outcomes. Prior studies have shown co-activation of the VS and CBL during various motor learning paradigms (*28*, *29*, *45*, *53*, *54*). Specifically, Tzvi et al., 2015 observed increased inhibition between the cerebellum and putamen in a motor learning task during pre- and post-sleep phases. Our work builds on these previous findings by proposing a potential mechanism by which the VS and CBL could communicate to integrate motor consequences with rewarding or non-rewarding outcomes.

Recent animal studies have identified functional and anatomical connections between the VS and CBL. Various studies have illustrated that the deep cerebellar nuclei, the output nuclei of the CBL, project to the VS through the ventral tegmental area (VTA) (*26*, *55*, *56*) or the intralaminar thalamus (*2*, *44*, *56*). Notably, electrical micro-stimulation of the deep cerebellar nuclei (DCN) elicited both excitatory and inhibitory modulation in the VS (*56*). In addition, food consumption in mice increases excitatory drive from the DCN to the VTA, suppressing VS activity through prolonged increases in basal dopamine concentrations, leading to decreased valuation of food and feelings of satiety (*55*). Thus, the CBL can flexibly modulate neuronal firing in the VS as a function of the motivational state. Our results advance this view by providing an additional functional basis by which the CBL could inhibit the VS following erroneous motor outcomes. While it is currently unclear whether there are direct anatomical projections from the VS to the CBL, anatomical tracing studies have found excitatory projections from the subthalamic nucleus (STN), a major output region of the basal ganglia, to the CBL (*3*, *44*). The STN is part of the indirect pathway in the BG and is thought to suppress movements (*57*). Following rewarding outcomes, the indirect pathway is suppressed to enable the reinforcement of the movement/action that led to the reward, which would result in the inhibition of the STN and, subsequently, the CBL (*58*). Thus, the STN could serve as a potential mediator that enables the VS to inform the CBL not to update the action that led to a rewarding outcome. Future studies could be designed to extend our causal models to include other subregions in the basal ganglia, such as the STN, to explore these connections further.

We observed that the VS-CBL connectivity following motor outcomes was amplified for individuals more willing to perform a motor task. Greater preferences for performing the motor task resulted in stronger inhibition from the VS to CBL following successful motor outcomes and from the CBL to VS following unsuccessful motor outcomes. Individuals with greater preferences for performing motor tasks could place more emphasis on processing the outcomes of their motor actions, which may be explained by a stronger need to dissociate between successful and unsuccessful motor outcomes. Our findings are consistent with previous work showing that subjective preferences and motivation can influence connectivity within the brain’s motor system. Previous work from our group has shown alterations in functional coupling between VS and motor cortical regions when individuals perform motor tasks for large prospective gains or losses (*12*). In addition, individuals with greater sensitivity to monetary losses over gains (i.e., loss aversion) exhibited increased motor cortical excitability for prospective gains in an incentive-based motor task (*36*). Furthermore, the CBL has been shown to encode movement parameters that are modulated by changes in motivation (*45*). In this context, our current results provide direct evidence that VS-CBL connectivity involves more than sensorimotor processing and is a potential means for the brain to integrate motoric outputs with motivational reward information.

We propose that the VS communicates differently with the sensorimotor and cognitive CBL. We found that the connection between the VS and the sensorimotor CBL primarily drove the inhibitory VS-CBL connectivity following motor outcomes. The sensorimotor cerebellum, consisting of the anterior cerebellar lobule (lobules I-V) and lobule VI, processes movement information and has shown functional connections with the sensorimotor network (*39*, *40*). Following an erroneous movement, the DCN, the output of the CBL, sends projections to the sensorimotor network to adjust motor commands (*17–19*). In addition to these projections, the sensorimotor CBL could also update the VS, ensuring that it does not reinforce erroneous motor actions. We observe a transition from excitation of the cognitive CBL by the VS to inhibition following successful motor outcomes for individuals with greater subjective preferences. Recent studies have shown that separate populations within granule and Purkinje cells of the posterior cerebellar regions (Lobule VI, Crus I & II) encode rewarding or unrewarding outcomes following a motor action (*23*, *46*, *59*). Wagner et al., 2017 have shown in rodents that throughout learning, there is an increase in the proportion of granule cells that respond to unrewarded outcomes while there is a decrease in the proportion of granule cells that respond to rewarded outcomes (*59*). As the granule cells comprise the input layer of the CBL, the VS may target distinct populations of granule cells, thereby enabling them to encode reward information. Purkinje cells, which receive input from granule cells have also shown similar dissociation between encoding rewarding and unrewarding outcomes (*23*, *46*), suggesting that a hierarchical organization could exist within the CBL that could be influenced by inputs from the VS. Importantly, how the VS interacts with these distinct populations within the cognitive cerebellum could explain interindividual differences in subjective preferences that we observe in our participants.

Our results implicate VS-CBL connectivity as a critical interface that ties motoric outputs with reward information and begins to shed light on potential mechanisms by which motoric and motivational deficits may be impacted in various patient groups. Patients with Parkinson’s disease (PD) have degeneration of dopaminergic neurons in the substantia nigra pars compacta (SNpc), and the resulting depletion of dopamine in the striatum leads to changes in basal ganglia activity and various motor deficits (*60*). It has been suggested that changes in basal ganglia activity lead to cerebellar hyperactivity in PD (*61–64*). Moreover, deep brain stimulation (DBS) and inhibiting STN output (STN-DBS) reduce cerebellar hyperactivity and relieve motor symptoms in PD patients (*61*, *65*, *66*). However, STN-DBS has also been shown to increase impulsivity, anhedonia, apathy, and other motivation-related deficits (*67*, *68*), consistent with the idea that BG/cerebellar connectivity is critical not just for sensorimotor control but also for motivational processes such as reward processing. In addition, impulsivity is observed in patients with obsessive-compulsive disorder (OCD) and has been associated with increased whole-brain connectivity between the striatum and cerebellar cortex (*69*). Our results further support these findings by suggesting that within healthy participants, the degree to which the VS interacts with cognitive cerebellar domains such as Crus I & II can be explained by an individual’s motivational state. Overall, the functional basis through which the BG and CBL take turns initiating communication with one another could prove crucial for understanding the motoric and motivational changes in neuropsychiatric and healthy populations.

## Materials and Methods

### Experimental setup

The game environment and behavioral data were presented and recorded using custom MATLAB scripts (http://www.mathworks.com) with the PsychToolBox library (Brainard, 1997). A projector placed at the back of the room displayed visual stimuli onto a rear-projection screen, which participants viewed through a 45-degree inclined mirror positioned above their heads.

For recording responses during the motor and behavioral choice tasks, participants used an MRI-compatible response box (Cedrus RB-830, Cedrus Corp., San Pedro, CA) with two horizontally aligned buttons, held in their right hand.

### Experimental Procedures

#### Participants

All participants were right-handed and were prescreened to exclude those with any history of neurological or psychiatric illnesses. This study was approved by the Institutional Review Board at the Johns Hopkins School of Medicine, and all participants provided informed consent.

Thirty participants initially participated in the experiment. Two subjects displayed significant motion inside the scanner (exceeding 3 mm a given axis) and were excluded from the group level analysis.

The final analysis included N = 28 participants in total (mean age, 24.9 years; age range, 20- 37 years; 16 females)

#### Motor Task

Prior to the experiment, participants were informed that they would receive a fixed show-up fee of $50. It was made clear that this fee did not depend on performance or behavior over the course of the experiment. The entire experiment occurred inside the fMRI scanner. However, participants were only scanned during the testing phase.

Visually, the task environment consisted of a horizontal line representing the floor and a vertical line with a middle gap representing a passage through the wall. The goal of the task was to control a cursor (red square) to jump through the target (i.e., passage in the wall). Hitting any part of the vertical wall during the jump resulted in failure. Each trial was initiated with a red square appearing to the left of wall and automatically moving horizontally to the right. Pressing the right button on the response box caused the cursor to jump, and letting go of the button caused the cursor to fall to the floor. As a result, the height of the jump was determined by the duration of the button press. To successfully complete the task, participants had to learn the precise timing and duration of the button press. The vertical trajectory of the cursor during the button press was determined using a sigmoid function to simulate acceleration induced by gravity during the jump.

The experiment began with participants completing 50 training trials under a fixed target size to become accustomed to the game environment and the button controls. The thresholding phase of the experiment was followed immediately after the training trials had been completed. The thresholding phase mirrored the training trials, except that every trial had varying target sizes on the wall, which affected the difficulty of completing the jump. There were 8 difficulty levels, with target sizes ranging from 26.5 to 52.4 mm (100 – 198 pixels). Each difficulty was presented ten times, which resulted in 80 trials in total for the thresholding phase.

Immediately after the thresholding phase was completed, a Weibull cumulative distribution function was used to model each participant’s performance and generate three difficulties (Easy, Medium, Hard), which would be used as the target sizes for the testing phase. Specifically, the psychometric model consisted of the following equation:

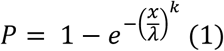

P is the mean success rate for trials with target size x, λ and k being the scale and shape parameters, respectively. The Levenberg-Marquardt nonlinear least squares algorithm was used to estimate the λ and k parameters. Based on the fitted model, the three target sizes for the main experiment were obtained by setting P equal to 0.9, 0.6 and 0.3, allowing difficulty levels to be controlled across participants.

During the testing phase, participants were scanned with fMRI while controlling the cursor through the target for potential reward. The testing phase differed from the training and thresholding phases in that following a successful jump, there was a probability of the floor breaking, which resulted in the cursor falling off the game environment. The thickness of the floor line indicated the probability of the floor breaking. A trial was considered successful only when the floor did not break. In addition, participants were presented with information regarding the probability of the floor remaining unbroken, which was either 30%, 60% or 90%, and the target size, which would be one of the three target sizes obtained from the thresholding phase.

At the start of each trial, information about the target size and floor thickness was presented sequentially in a pseudorandom order (**Figure 1A**) (e.g. medium difficulty presented, followed by floor presentation with 30% of not breaking, each with jittered duration of 2–4 s). Then, participants performed the motor task (∼1 s duration) and were shown the outcome of their performance (jittered duration, 2-4 s). If the task was unsuccessful, the word “Failure” appeared below the floor with a red color. On the other hand, successfully completing the motor task led to the outcome of the floor being revealed (jittered duration, 2-4 s). The green “Success” text appeared on the screen if the floor did not break. If the floor did break, the right side of the floor environment disappeared, the cursor fell off the game environment, and the red “Failure” text appeared on the screen. Accounting for the order of floor/task difficulty presentation, there were 18 different trial combinations. Each of the 5 sessions had 36 trials (∼10 minutes), where each trial combination was shown twice in a randomized order, which resulted in a total of 180 trials. Participants were informed that at the end of the experiment, one of the trials would be chosen randomly and would be paid $25 if the trial was a success.

#### Decision-Making Task

This task was performed after completing the thresholding phase and before starting the testing phase. Participants made choices (4 s limit) to either perform a motor task of varying difficulty (i.e., target size) or accept probabilistic outcomes that were not associated with task performance (i.e., monetary gamble) (**Figure 4A**). The target sizes were sampled from those generated in the thresholding phase (Easy – 30%, Medium – 60%, and Hard-90% difficulty) while the monetary gamble had success probability ranging from 0.10 to 0.90 (in increments of 0.10). There were 135 trials where each of the 27 different pairs of decision trials were shown five times in randomized order. Participants were told that at the end of the experiment, one of the trials would be chosen randomly, and they would then have to play out the trial depending on their response. If the participant chose the task, they received $25 upon successfully completing the motor task. The success criteria were the same as the training and thresholding phases. If the participant accepted the probabilistic outcome, they received $25 when the monetary gamble was successful. The monetary gamble was simulated from a Bernoulli distribution with a probability of success, which was set according to the trial.

#### MRI protocol

The MRI scanning sessions were conducted on a 3 Tesla Philips Achieva Quasar X-series MRI scanner equipped with a radio frequency coil. High-resolution structural images were acquired using a standard MPRAGE pulse sequence, covering the entire brain at a resolution of 1 mm × 1 mm × 1 mm. Functional images were collected at a 30° angle from the anterior commissure-posterior commissure (AC-PC) axis, with 48 slices acquired at a resolution of 1.87 mm × 1.87 mm × 2 mm to ensure full brain coverage. An echo-planar imaging (FE EPI) pulse sequence was applied (TR = 2800 ms, TE = 30 ms, FOV = 240, flip angle = 70°).

As physiological variables have been shown to introduce confounds in fMRI data (*50*, *70–72*), cardiac and respiration rates were obtained during the functional scans using a peripheral pulse unit and a respiratory belt with a sampling rate of 496 Hz (wireless). The physiological recordings were then processed using model-based noise correction via the PhysIO Toolbox (*72*), which provided an output of multiple regressor matrix containing explanatory components of physiological noise. To account for significant head motion, scans with root mean square of translational or rotational head motion exceeding 0.5 mm/deg were included as nuisance regressors in subsequent analyses.

### Data Analysis

#### Behavioral Performance Analysis

To measure changes in performance across the training session, the training trials (n = 50) for each participant were binned into 5 blocks where each block consisted of ten trials. The mean success rate for each block was then obtained for each participant. A one-way ANOVA was used to determine whether there was significant difference in success rate across the training blocks. To measure changes in performance across the testing session, the mean success rate in the motor task for each imaging session was obtained for each participant. A one-way ANOVA was used to determine whether there was significant difference in success rate across the sessions. Individual p-values for each comparison for both testing and training sessions were corrected for multiple comparisons using Tukey’s honestly significant difference procedure. To identify variables that influenced performance in the motor task, a generalized linear mixed-effects model was used to predict trial success (binary variable indicating 1 if success, 0 otherwise in the motor task) as a function of target size, the probability of floor remaining unbroken and session number. The distribution of trial success was specified as a binomial distribution and the model was estimated via maximum likelihood through Laplace approximation.

#### Image Analysis

The SPM12 software package was used to analyze the fMRI Data (Wellcome Trust Centre for Neuroimaging, UCL, London, UK). A slice-timing correction was applied to the functional images to adjust for the fact that different slices within each image were acquired at slightly different time points. Images were corrected for participant motion by registering all images to the first image, spatially transformed to match a standard echo-planar imaging template brain, and smoothed using a 3D Gaussian kernel (5 mm FWHM) to account for anatomical differences between participants. The final resolution of the preprocessed images had voxel dimensions of 2.00 mm x 2.00mm x 2.00mm. This set of data was then analyzed statistically.

### General Linear Model

The general linear model (GLM) was used to generate voxel-wise statistical parametric maps from the fMRI data. We created GLMs that included conditions relevant to the skilled motor task shown in Figure 1A. The outline of the GLM that was constructed for each participant is shown below:

Model

1. The presentation of the target size (Task Presentation, 2-4 seconds)
2. The presentation of the floor probability (Floor Presentation, 2-4 seconds)
3. Motor task (Task Performance, 1 second)
4. Successful Motor Outcomes (Motor Success, 2-4 seconds)
5. Unsuccessful Motor Outcomes (Motor Failure, 2-4 seconds)
6. Successful Probabilistic Outcomes (2-4 seconds)
7. Unsuccessful Probabilistic Outcomes (2-4 seconds)

Separating conditions for successful and failed task performance outcomes allowed us to contrast the two conditions as shown in Figure 2. All conditions were modeled as blocks except for the motor task, which was modeled as event regressors. In addition to the main effects, each 1^st^ level GLM model shown above contained a multiple-regressor-matrix obtained from the PhysIO Toolbox, which contained regressors for the movement, cardiac, respiratory and motion outlier nuisance regressors.

#### Region of Interest Analysis and Statistical Tests

To create a ROI for the VS, we identified peak coordinates from a meta-analysis (*73*) which identified brain regions involved with processing of monetary gains and losses (left VS MNI coordinates (x,y,z) = [12,10,-4]; right VS MNI coordinates (x,y,z) = [12,10,-4]). The bilateral VS mask was thus obtained by first creating 6 mm spheres around the left and right VS coordinates and combining the spheres using MarsBaR (*74*). To generate an ROI for the CBL, we utilized the leave-one-subject-out (LOSO) approach (*75*). We defined CBL ROIs by taking a conjunction between an anatomical CBL mask and a group-level Motor Failure / Motor Success contrast (Figure 2C) from all but one participant set at p < 0.001. The overlap between these two masks would then be used as the CBL ROI for the left-out participant for the correlation and the first DCM analysis.

To obtain separate ROIs for the Mtr. and Cog. CBL, we repeated the same LOSO approach but specified an anatomical Mtr. CBL mask (including lobules I-VI) and Cog. CBL mask (Crus I & II) when taking the conjunction between the Motor Failure / Motor Success contrast.

To display whole-brain contrasts and report CBL activations, we used a threshold of p < 0.001 with a 20-voxel extent threshold. The Cerebellar activations are displayed on a cerebellar flat map using the SUIT toolbox (*76*). Statistical inference was performed within the SPM framework using family-wise error correction for the whole brain. To report the correlations between the VS and CBL parameter estimates, we report Kendall’s rank correlation due to the non-normal distribution of VS and CBL parameter estimates.

#### Dynamic Casual Modeling (DCM)

To investigate the connectivity between the VS and CBL underlying task performance outcomes, we constructed deterministic bilinear DCMs. DCM utilizes forward modeling to represent how experimental manipulations lead to change in neural activity through a bilinear differential equation in the form of:

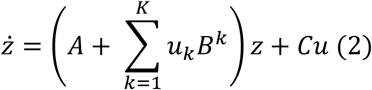

where *ż* represents the change in neural activity per unit time (i.e., derivative of neural activity for each region), u represents experimental conditions (i.e., successful or failed task performance outcomes), A is a matrix that defines the intrinsic coupling (average effective connectivity) between regions, B is a matrix specifying the modulation of effective connectivity due to experimental condition k = 1…K, and C is a matrix encoding the sensitivity of each region to driving input from experimental condition k = 1…K. Each connection thus represents a change in neural activity due to directed signals (A, C matrix), or the modulation of the directed connection (B matrix).

We followed the procedures described in Zeidman et al. (*32*, *33*) for specifying DCM across participants. For our first DCM analysis, we assigned input nodes as the VS and CBL, and both regions were set to be driven by experimental conditions including both successful and failed motor outcomes (i.e., C matrix). We enabled full bidirectional connections between the VS and CBL (i.e., A matrix). For the second DCM analysis, we assigned input nodes as the VS, Mtr. CBL, and the Cog. CBL. All three regions were set to be driven by successful and failed task performance outcomes (i.e., C matrix). We enabled full bidirectional connections (i.e., A matrix) between all regions except for connections between the Mtr. and Cog. CBL. For both DCM analyses, we were specifically interested in parameters modeling the modulation in the VS-CBL connectivity following successful and failed task performance (i.e., B matrix).

#### Parametric Empirical Bayes (PEB)

The PEB approach uses hierarchical Bayesian modeling where each parameter (i.e., the estimate of connection strength) is treated as a random effect. All participants are assumed to have the same model architecture but varying strengths of connections within the group (*33*). Thus, one full DCM is specified per participant, enabling all connections, and a series of candidate models to be specified that test different connectivity hypotheses. The model evidence (Free energy) is then computed for each model, which is the trade-off between the accuracy (i.e., ability to predict observed BOLD activity) and complexity (KL-divergence between estimated parameter and the original priors) (*32*, *77*). We can then perform a Bayesian model comparison where the model evidence is compared across all models. This process results in a posterior probability, corresponding to how much each model contributes to the overall model evidence and, thus, provides the best explanation of the observed data. To obtain PEB estimates of individual parameters (i.e., estimation of connection strength), we performed Bayesian Model Average (BMA) where the average of the parameters form different models are weighted by the models’ posterior probabilities (*34*, *35*). We present and interpret the results of PEB parameters with positive (p > 0.73) and very strong (p > 0.99) Bayesian evidence.

We followed the procedures described in Zeidman et al. (*40*) for performing PEB estimation. For our first DCM analysis, the PEB design matrix included 1) a constant term, 2) mean-centered mean performance, and 3) mean-centered subjective preference. Due to the mean-centering, the estimated parameters represent the mean coupling strength between the VS and CBL following task performance and the additive effect of the covariates (mean performance & subjective preferences) on this common effect. For the second DCM analysis, our PEB design matrix included only the constant and mean-centered subjective preference as regressors, as we were interested in supplementing the findings observed in our first DCM analysis. Following PEB estimation, we utilized leave-one-out (LOO) cross-validation using the observed effects of interest (i.e., modulation from VS to CBL following successful task performance) to estimate the out-of-sample subjective preference for each participant. The LOO approach provides a statistically robust approach to assess the validity of associations (*33*). We report the Pearson correlation coefficient (R) and the p-value for right-tailed correlation between the model estimated and observed subjective preferences.

## ACKNOWLEDGEMENTS

This work was supported by the Eunice Kennedy Shriver National Institute of Child Health & Human Development of the National Institutes of Health under Award Number R01HD097619 and the National Institutes of Mental Health under Award Numbers R56MH113627 and R01MH119086 to V.S.C; and the National Institute on Deafness and Other Communication Disorders R01DC018061 to K.E.C.

