## Supplementary Figures for "Striatal and cerebellar interactions during reward-based motor performance"

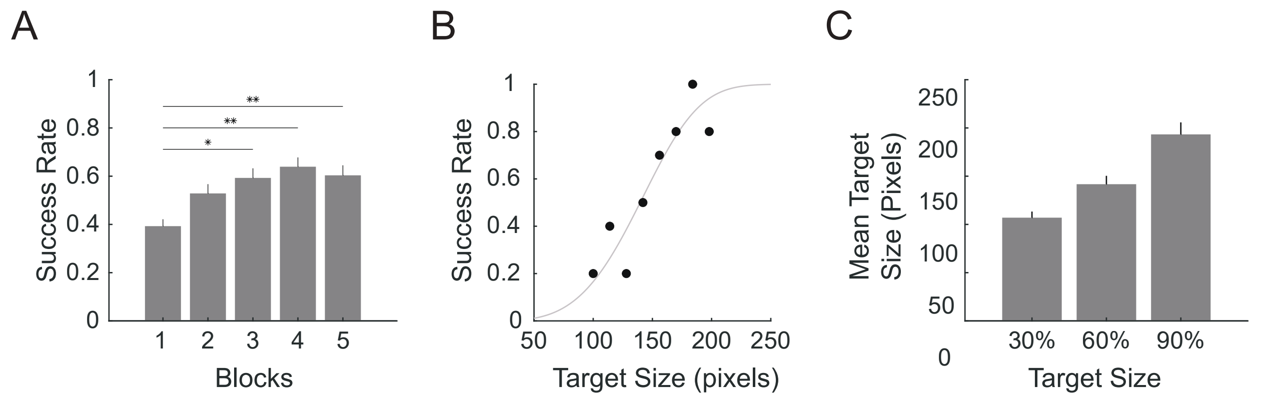


### Supplementary Figure 1: Analysis of participants' performance during the training and thresholding phases

**(A)** Performance in the motor task improved and stabilized throughout the training session. Performance was measured by dividing training trials (n = 50) into 5 blocks and computing mean success rate across trials in each block (n = 10). There was a significant difference in performance across sessions ($F(4,135) = 6.65$, $p < 0.0001$). (*$p < 0.005$, ** $p < 0.001$ corrected for multiple comparisons using Tukey's honestly d significant difference procedure). Error bars denote SEM.

**(B)** Exemplary participant's performance across different target sizes (pixels). A Weibull cumulative distribution function (gray curve) was used to model participants' performance and estimate three target sizes where the expected success rate would be 30%, 60%, and 90% for the easy, medium, and hard difficulties respectively.

**(C)** Mean target size (pixels) for the three difficulties for all participants. Error bars denote SEM.

A

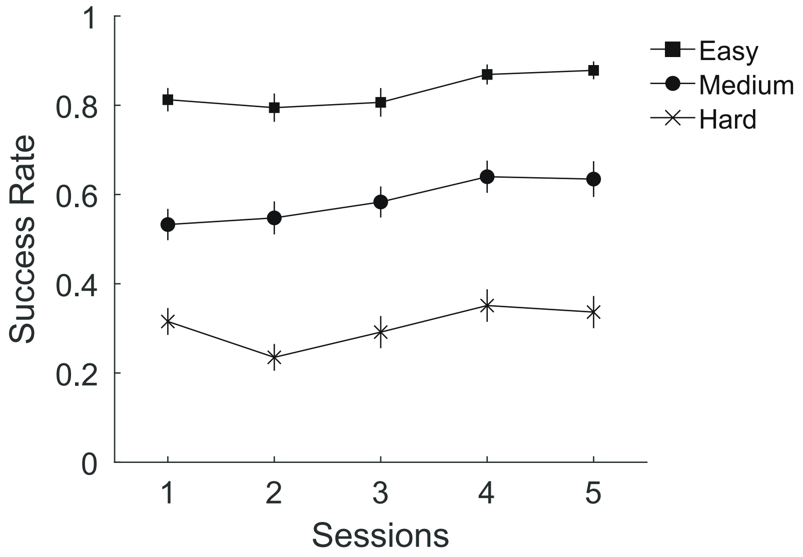


B

| Fixed Effects Coefficients | Estimate | Standard Error | T | Degrees of Freedom | P Value | Confidence Interval |
|---|---|---|---|---|---|---|
| Intercept | -0.030711 | 0.031757 | -0.96708 | 1202 | 0.3337 | [-0.093, 0.032] |
| Target Size | 0.26306 | 0.0084267 | 31.218 | 1202 | < 0.0001 | [0.247, 0.280] |
| P (Floor) | 0.0052861 | 0.0084267 | 0.6273 | 1202 | 0.53058 | [-0.011, 0.022] |
| Session | 0.02315 | 0.0049979 | 4.632 | 4961 | < 0.0001 | [0.013, 0.033] |

**Supplementary Figure 2: Analysis of participants' performance during the motor task**

**(A)** Participants' mean success rate in the motor task throughout the five experimental sessions for each target size.

**(B)** Table summarizing the mixed-effects GLM modeling participants' performance in the motor task as a function of the target size, probability of floor remaining stable (P (floor)) and session.

| Cerebellar region | Peak MNI coordinates (mm) | | | Peak t value | Cluster size (Voxels) |
|---|---|---|---|---|---|
| | x | y | z | | |
| Lobules 4/5 R | 24 | -36 | -22 | 7.83 * | 579 |
| Lobules 4/5 R | 28 | -46 | -22 | 7.38 * | |
| Lobule 6 R | 30 | -80 | -20 | 6.46 * | |
| Crus I L | -40 | -70 | -22 | 6.66 * | 596 |
| Crus II L | -26 | -76 | -44 | 6.27 | |
| Lobule 6 L | -32 | -48 | -22 | 6.14 | |
| Lobule 4/5 L | -26 | -34 | -39 | 5.23 | 25 |

**Supplementary Table 1: Cerebellar activations for the contrast Failure - Success at time of motor outcome**

* Cerebellar Regions that survive whole-brain correction for multiple comparisons (FWE) at $p < 0.05$. Laterality - right (R); left (L).

A

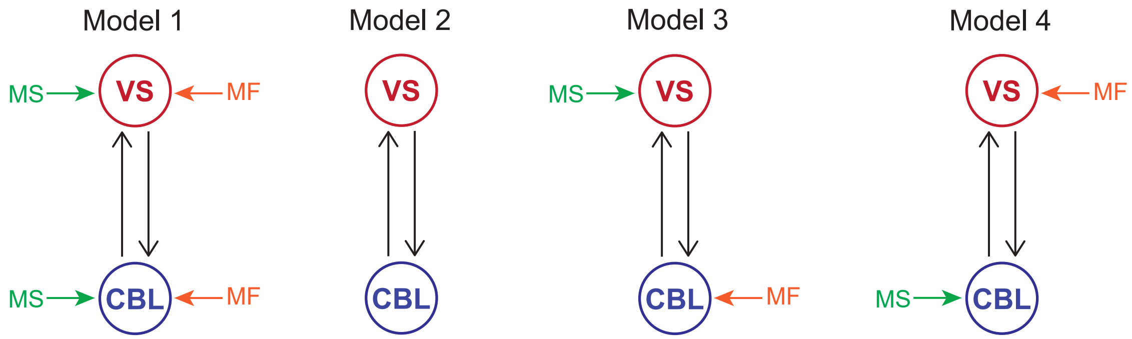


B

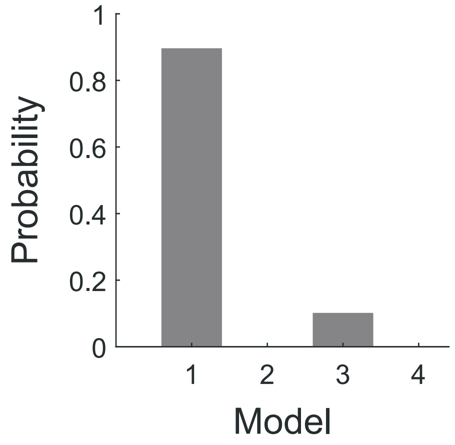


C

| Driving Input | Target | Connectivity (Hz) |
|---|---|---|
| Motor Success | VS | 0.067 ** |
| Motor Success | CBL | 0.070 * |
| Motor Failure | VS | 0.167 * |
| Motor Failure | CBL | 0.187 ** |

**Supplementary Figure 3: Successful and unsuccessful motor outcomes modeled as driving inputs to the VS and CBL.**

**(A)** We proposed four models that explain how motor outcomes could potentially drive activity in the VS and CBL. MS = Motor Success, MF = Motor Failure.

**(B)** A Bayesian model comparison showed that model 1 was the winning model (90%), followed by model 3 (10%) for describing how successful and failed motor outcomes drives VS and CBL activity.

**(C)** BMA of PEB parameters highlighting how motor outcomes drive VS and CBL activity. ** PEB parameters showing very strong positive Bayesian evidence (posterior probability > 0.99). * PEB parameters showing positive Bayesian evidence (posterior probability > 0.73).

A

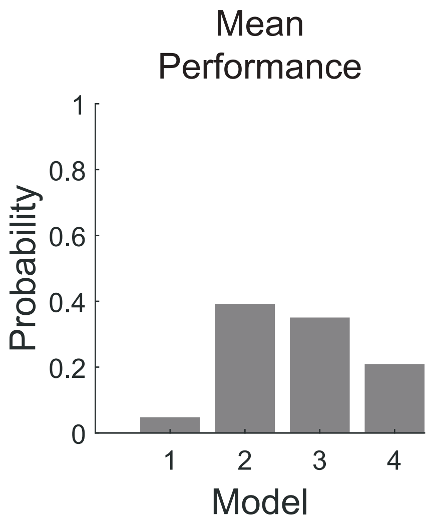


B

| Driving Input | Source | Target | Connectivity (Hz) | | |
|---|---|---|---|---|---|
| | | | **Commonalities** | **Subjective Preference** | **Mean Performance** |
| Motor Success | VS | CBL | -0.048 ** | -11.294 ** | 0.40 |
| Motor Success | CBL | VS | -0.024 | -0.374 | 0.26 |
| Motor Failure | VS | CBL | 0.017 | 0.001 | 0.26 |
| Motor Failure | CBL | VS | -1.338 ** | -2.190 ** | 0.40 |

### Supplementary Figure 4: BMA of PEB parameters for first DCM analysis.

**(A)** A Bayesian model comparison showed that models 2 (44%), 3 (38%) and 4 (21%) best described the effect of participants' mean performance on their VS-CBL connectivity.

**(B)** Modulation effect detailing all connections in the full model (Model 1) for the first DCM analysis (**Figures 3-4**). Commonalities refers to the mean strength of the connection (Hz) between the source and target regions. Negative values correspond to inhibition whereas positive values correspond to excitation. The subjective preference and mean performance were two covariates that were included as second-level effects. Subjective preference corresponded to participants' tendency to choose the motor task over the probabilistic coin flip for potential rewards in the choice task (See **Figure 4A**). Mean performance corrsponded to participants' overall success rate in the motor task. ** PEB parameters showing very strong positive Bayesian evidence (posterior probability > 0.99). * PEB parameters showing positive Bayesian evidence (posterior probability > 0.73).

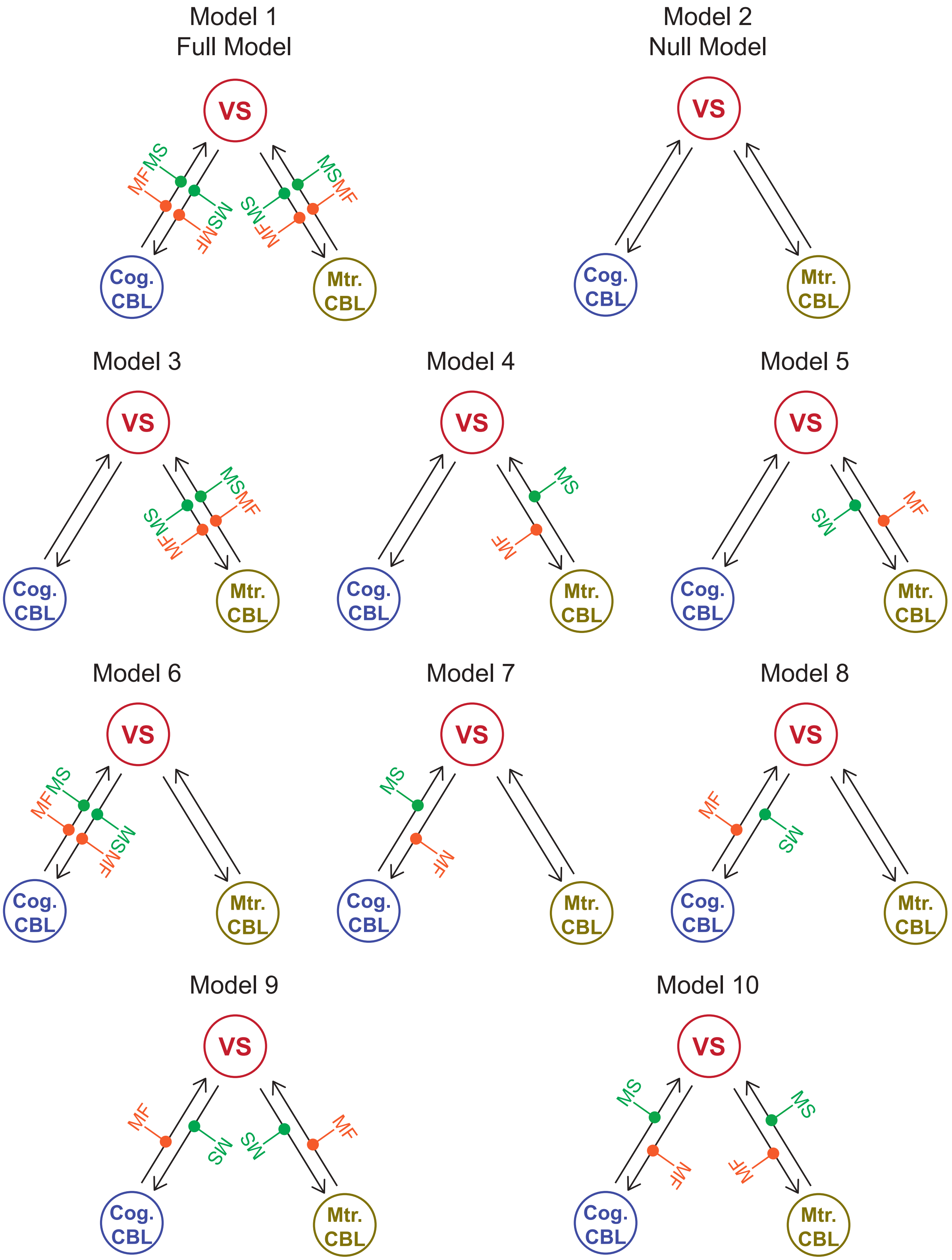

Model 1
Full Model
Model 2
Null Model
Model 3
Model 4
Model 5
Model 6
Model 7
Model 8
Model 9
Model 10
VS
Cog. CBL
Mtr. CBL
MS
MF

### Supplementary Figure 5: Visualization of all models tested in the second DCM analysis

For each model, successful (MS) and failed (MF) motor outcomes (Factor 1) outcomes modulated the connectivity between the VS and the Mtr. / Cog. CBL (Factor 2) depending on the hypotheses specified in Figure 5B. Along with a 'null' model which had no modulation, there were a total of 3 x 3 + 1 = 10 models specified.

| Driving Input | Source | Target | Connectivity (Hz) | |
|---|---|---|---|---|
| | | | Commonalities | Subjective Preference |
| Motor Success | VS | Ant. CBL | -0.662 ** | -0.394 |
| Motor Success | VS | Post. CBL | 0.260 * | -4.855 * |
| Motor Success | Ant. CBL | VS | 0.172 | -2.849 |
| Motor Success | Post. CBL | VS | -0.223 | 2.131 * |
| Motor Failure | VS | Ant. CBL | 0.088 | 2.311 |
| Motor Failure | VS | Post. CBL | -0.150 | 2.999 * |
| Motor Failure | Ant. CBL | VS | -0.825 ** | 0.687 |
| Motor Failure | Post. CBL | VS | -0.425 * | -5.091 * |

### Supplementary Table 2: BMA of PEB parameters for second DCM analysis.

Modulation effect detailing all connections in the full model (Model 1) for the second DCM analysis (Figure 5). ** PEB parameters showing very strong positive Bayesian evidence (posterior probability > 0.99). * PEB parameters showing positive Bayesian evidence (posterior probability > 0.73).

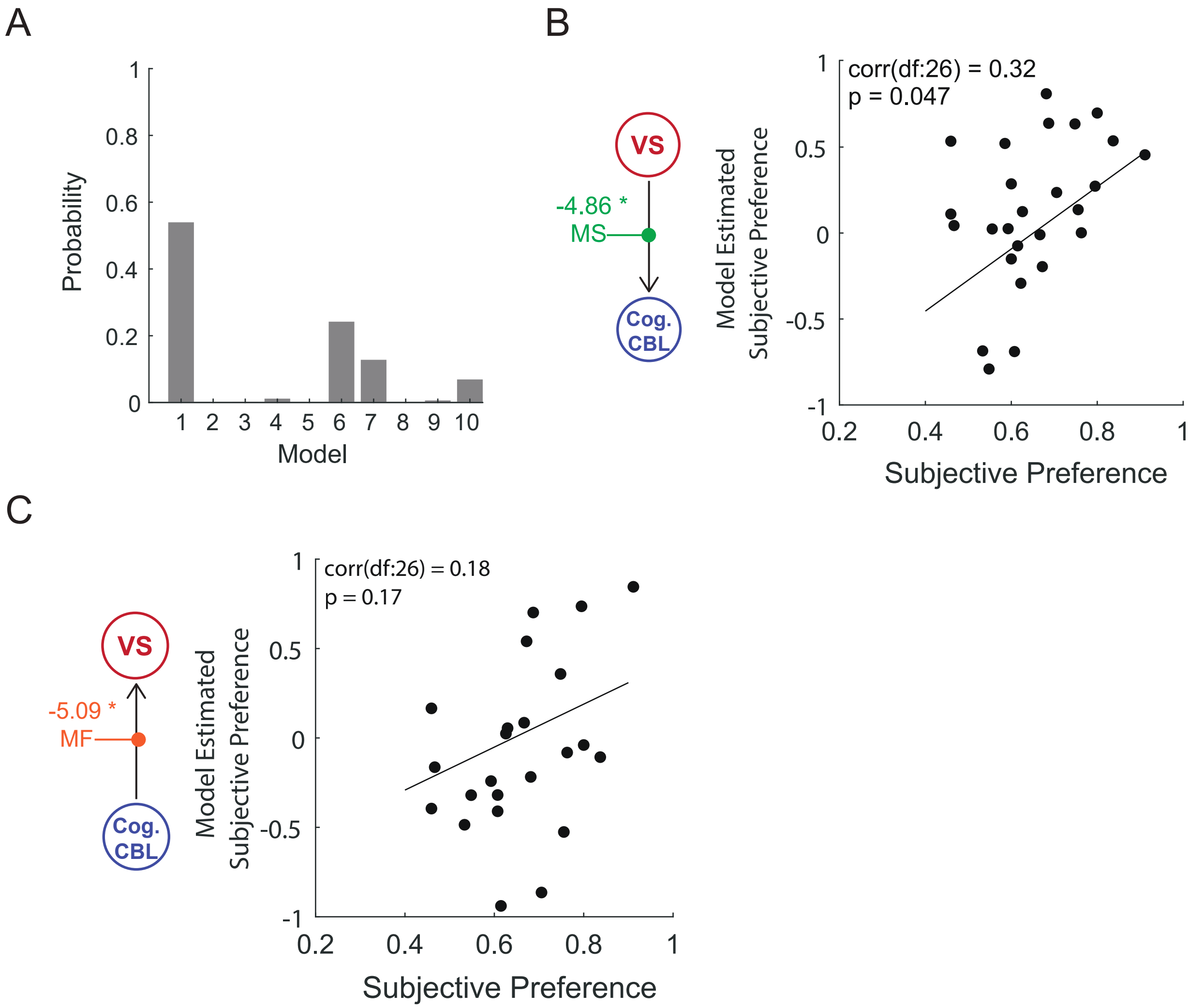


**Supplementary Figure 6: Model comparison and cross-validation estimates for how participants' subjective preference influences VS - Cog. CBL connectivity.**

**(A)** A Bayesian model comparison showed that model 1 (54%), followed by models 6 (24%) and 7 (13%) best described the effect that participants' subjective preference has on their VS-Mtr. CBL and VS- Cog. CBL connectivity.

**(B)** Out-of-sample estimation of participants' subjective preference using the degree of inhibition of the Cog. CBL by the VS following successful motor outcomes (PEB parameter, Left).

**(C)** Out-of-sample estimation of participants' subjective preference using the degree of inhibition of the VS by the Cog. CBL following unsuccessful motor outcomes (PEB parameter, Left).
